## Supplementary Information for "Hierarchical Morphogenesis of Swallowtail Butterfly Wing Scale Nanostructures"

### Supplementary Methods:

One-sided ($H_a$: treatment < control) pairwise *t*-tests with default settings (pooled standard deviations, Holm method to correct for multiple comparisons) were performed in the *R* statistical environment (version 4.1.2) running on macOS Monterey (ver. 12.6.5).

**Figure S1.** Diversity of adult wing scale nanostructure in Papilionidae shown in comparison to (R) a dorsal forewing (DFW) black scale of *Hypolimnas bolina* (Nymphalidae) with regular rectilinear cross-ribs. *P. arcas*: (A) male DFW green cover, (B) male DFW black, (C) female DFW yellow cover, (D) female dorsal hindwing (DHW) red cover; *P. eurimedes*: (E) male DFW green cover, (F) proximal part of male DFW black; *P. nireus*: (G, H) male DFW blue cover; *P. memnon*: (I) DFW navy, (J) Ventral hindwing (VHW) black, (K) VHW red; *P. palinurus*: (L) DHW green cover, (M) DFW green cover, (N) Ventral forewing (VFW) dark gray, (O) VHW blue; *Graphium agamemnon*: (P) light pink, (Q) dark pink; (R) DFW black; *Hypolimnas bolina*. All scale bars – 1μm except for (L) – 5μm. Abbreviations: r – ridges, mr – microribs, cr – crossribs, hc – honeycombs, t – trabeculae, lpc – lumen photonic crystal, d – dimples.

**Figure S2.** Early development of dorsal forewing green cover scales in pupal male *P. eurimedes* acquired with 60x confocal microscope. AF-555 WGA (green) stains plasma membrane and AF-647 phalloidin (red) stains F-actin. (A-A'') 38% development: Young scales extend out from the wing epithelium. F-actin filaments are beginning to organize into bundles. (B-B'') 43% development: F-actin bundles span the scale length along the proximal-distal axis. Plasma membrane of scale cells begins to pleat. (C-C'') 48% development: Ridge-like projections are seen in between adjacent F-actin bundles with WGA faintly staining crossribs. All scale bars – 5μm.

**Figure S3.** Yellow dorsal forewing cover scales of pupal female *P. eurimedes* at 48% development acquired with 100x confocal microscope. AF-555 WGA (green) stains chitin and Cellmask (red) stains plasma membrane. Scales are relatively young with a long and narrow appearance. (B-B'') Close up views of A-A''. Ridges seen on the surface of scale cells are indistinct while the membrane has a mottled appearance. (C-C'') *xz* cross-sections of the scale at location marked with grey line in A'' showing chitin is deposited on the cell membrane, as expected. Scale bars (A-A'') – 5μm, (B-B'' and C-C'') – 2μm.

**Figure S4.** Yellow dorsal forewing cover scales of pupal female *P. eurimedes* at 52% development acquired with 100x confocal microscope. AF-555 WGA (green) stains chitin and Cellmask (red) stains plasma membrane. Cellmask staining reveals hollow anastomosing vein-like crossribs on the plasma membrane in between ridges that will serve as boundaries for cuticle accretion. (B-B'') Close up views of A-A'' showing chitin is deposited within the hollow membranous outlines. (C-C'') *xz* cross-sections of the scale at location marked with grey line in A''. Most of the chitin deposition is on the scale surface. Yellow ROI in C'' corresponds to that in B''. Scale bars (A-A'') – 5µm, (B-B'' and C-C'') – 2µm.

**Figure S5.** Yellow dorsal forewing cover scales of pupal female *P. eurimedes* at 62% development acquired with 100x confocal microscope. AF-555 WGA (green) stains chitin and Cellmask (red) stains plasma membrane. As scales mature, more cuticle is deposited on rows of disks bounded by the membraneous crossribs as compared to the rest of the scale cell surface. (B-B'') Close up views of A-A''. (C-C'') *xz* cross-sections of the scale at location marked with grey line in A'' reveal the planar aspect of the cuticular disks, for the honeycomb lattice has not formed yet. Yellow ROI in C'' corresponds to that in B''. Scale bars (A-A'') – 5µm, (B-B'' and C-C'') – 2µm.

**Figure S6.** Black dorsal forewing scales of pupal male *P. eurimedes* at approximately 62% development acquired with 100x confocal microscope. AF-555 WGA (green) stains chitin and Cellmask (red) stains plasma membrane. (B-B'') Close up views of A-A''. Cuticular disks are clearly seen in between ridges and within the hollow outlines (crossribs) defined by the plasma membrane. (C-C'') *xz* cross-sections of the scale at location marked with grey line in A'' again shows the cuticular disks are flat and bulk of the cuticle deposition is in the cuticular disks. Yellow ROI in C'' corresponds to that in B''. Scale bars (A-A'') – 5µm, (B-B'' and C-C'') – 2µm.

**Figure S7.** Green dorsal forewing cover scales of pupal male *P. eurimedes* at 57% development acquired with super-resolution lattice SIM. (A-A'') AF-555 WGA (green) stains double rows of cuticular disks along the upper lamina. F-actin bundles as stained by AF-647 phalloidin are seen disintegrating into shorter fibrils that are distributed around the cuticular disks. The scale is still soft (ridges are not fully sclerotized) at this point, given the irregular, dimpled appearance. (B-B'') Close up views of A-A''. Insets correspond to the regions of interest (ROI) marked in yellow shown with a 3D aspect. (C-C'') *xz* cross-sections of the scale at locations marked with grey lines in A-A''. Yellow ROI correspond to those in B-B''. Scale bars (A-A'') – 5µm, (B-B'', C-C'' and insets) – 1µm.

**Figure S8.** Green dorsal forewing cover scales of pupal male *P. eurimedes* at 67% development acquired with super-resolution lattice SIM. AF-555 WGA (green) stains chitin and AF-647 phalloidin (red) stains F-actin. (A-A'') The cuticular disks are more regularly arranged, as the scale cells expand and flatten. (B-B'') Close-up views of (A-A'') show the disintegrating F-actin bundles have reorganized into a mesh-like network and associate with the cuticular disks to a greater degree. The cuticular disks now appear elongated and slightly tubular in cross-sections. Insets correspond to the ROI marked in yellow shown with a 3D aspect. (C-C'') *xz* cross-sections of the scale at locations marked with grey lines in A-A''. Yellow ROI correspond to those in B-B''. The cuticular disks are located along the upper lamina surface whereas F-actin structures extend further into the lumen. Scale bars (A-A'') – 5µm, (B-B'', C-C'' and insets) – 1µm.

**Figure S9.** Black dorsal forewing scales of pupal male *P. eurimedes* at 76% development acquired with super-resolution lattice SIM. AF-555 WGA (green) stains chitin and AF-647 phalloidin (red) stains F-actin. (A-A'') Scale cells take on the overall appearance of adult black scale cells with finger-like projections on the distal tip. (B-B'') Close-up views of (A-A'') show F-actin has completely disintegrated. The reorganized F-actin enmeshes the cuticular disks, which have extruded into the lumen to become hollow tubes at this stage. Insets correspond to the ROI marked in yellow shown with a 3D aspect. (C-C'') *xz* cross-sections of the scale at locations marked with grey lines in A-A''. Yellow ROI correspond to those in B-B'' and shows higher degree of association as compared to earlier stages. Scale bars (A-A'') – 5μm, (B-B'', C-C'' and insets) – 1μm.

**Figure S10.** Yellow dorsal forewing cover scales of pupal female *P. eurimedes* at approximately 52% development acquired with 100x confocal microscope. AF-594 anti-Arp2 (green) stains Arp2/3 complex and AF-647 phalloidin (red) stains F-actin. (A-A'') Arp2/3 complex shows a punctate, sparse distribution, while F-actin bundles are intact. (B-B'') Close-up views of (A-A''). (C-C'') *xz* cross-sections of the scale at locations marked with grey lines in A-A''. Yellow ROI correspond to those in B-B''. Scale bars (A-A'') – 5μm, (B-B'' and C-C'') – 2μm.

**Figure S11.** Yellow dorsal forewing cover scales of pupal female *P. eurimedes* at approximately 62% development acquired with 100x confocal microscope. 488nm channel shows cuticular autofluorescence from scales. AF-594 anti-Arp2 (green) stains Arp2/3 complex and AF-647 phalloidin (red) stains F-actin. (A-A''') F-actin bundles have disintegrated at this stage and re-organized into a reticulate network seen closely associating with a relatively higher concentration of punctate Arp2/3 complex. (B-B''') Close-up views of (A-A'''). (C-C''') *xz* cross-sections of the scale at locations marked with grey lines in A-A'''. Yellow ROI correspond to those in B-B'''.

Ridges are seen in the autofluorescence channel. Scale bars (A-A''') – 5µm, (B-B''' and C-C''') – 2µm.

**Figure S12.** Green dorsal forewing cover scales of pupal male *P. eurimedes* at approximately 76% development acquired with 100x confocal microscope. 488nm channel shows cuticular autofluorescence from scales. AF-594 anti-Arp2 (green) stains Arp2/3 complex and AF-647 phalloidin (red) stains F-actin. (B-B''') Close-up views of (A-A'''). (C-C''') *xz* cross-sections of the scale at locations marked with grey lines in A-A'''. Yellow ROI correspond to those in B-B'''. F-actin network shows a greater degree of reorganization. A large amount of the Arp2 antibody signal overlaps with cuticle autofluorescence, but the punctate pattern can still be discerned. Scale bars (A-A''') – 5µm, (B-B'' and C-C''') – 2µm

**Figure S13.** Negative controls for Arp2 antibody staining. Red dorsal hindwing cover scales from the same pupal male *P. eurimedes* from Fig. S12 (A-A'' at approximately 62% development), and Fig. S13 (C-C'' at approximately 76% development) acquired with 100x confocal microscope. 488nm channel shows cuticular autofluorescence from scales while AF594 antibody serves as a negative control for unspecific binding from the secondary antibody. Cuticle shows autofluorescence at higher wavelengths as scale matures. (B-B'', D-D'') *xz* cross-sections of the scale at locations marked with grey lines in A-A'' and C-C'' respectively. Scale bars (A-A'', C-C'') – 5µm, (B-B'', D-D'') – 2µm

**Figure S14.** Pharmacological disruption of Arp2/3 with CK-666 in *Parides* pupae. Violin plots depict the distribution of honeycomb lattice heights measured from multiple scales of a single emerged male individual either treated with CK-666 or DMSO (negative control) as pupae, along with representative SEM images of scale cross-sections used in measurements for *P. eurimedes* (A-C) and *P. iphidamas* (D-F). Developmental stage of the pupae at the time of injection are as marked. Results of pairwise *t*-tests (after corrections for multiple comparisons) are indicated as follows: * $P < 0.05$, ** $P < 0.01$, *** $P << 0.01$

**Figure S15.** Development of dorsal forewing black cover scales in pupal *P. polytes* acquired with 100x confocal microscope. AF-555 WGA (green) stains chitin and AF-647 phalloidin (red) stains F-actin. (A) Adult *P.polytes*. Credit: iDigBio YPM ENT 815256 (reproduced as CC0 1.0). (B) SEM top view and (C) cross-sectional image of an adult black scale showing irregular crossribs (white arrow). (D-D'') 50% development: Chitinous ridges developed in between adjacent F-actin bundles. (E-E'') *xz* cross-sections of the scale at location marked with grey line in A'' shows F-actin bundles along the periphery of scale cells, with thicker bundles at the lower surface of the cell, (F-F'') 60% development: WGA signal shows irregular crossrib pattern that resemble the final morphology in adult scales. (H-H'') 70% development: WGA signal is fuzzier, possibly due to the increased amount of lipids and waxes deposited in the cuticle. Irregular crossribs are still visible in between the ridges. (G-G'' and I-I'') *xz* cross-sections of the scale at location marked with grey lines in F'' and H'' respectively. Scale bars (B) – 2μm, (D-D'', F-F'' and H-H'') – 10μm, (E-E'', G-G'' and I-I'') – 5μm.

**Figure S16.** CK-666 inhibition does not affect F-actin structure in *P. polytes*. *P. polytes* pupae were injected with 100μM CK-666 at 50%, 60%, and 70%

development, and dissected at 80% development. Dorsal forewing cover scales are stained with AF-555 WGA (green) and AF-647 phalloidin (red) and acquired with 100x confocal microscope. Ridges and irregular crossribs developed across all treatment days. (B-B'', D-D'' and F-F'') *xz* cross-sections of the scale at locations marked with grey lines in A'', C'' and E'' respectively. All scale bars – 10μm.

**Figure S17.** SEM micrographs of *P. polytes* pupal scales injected with CK-666 at different developmental stages. Wing scales are from the same pupa corresponding to Fig. S16, but are different black scales. Irregular crossrib formation is not disrupted after CK-666 treatment. All scale bars – 1μm.

**Supplementary Movie 1.** Planar sections through the same dataset (57% development) as depicted in Fig. S8. AF-555 WGA (green) stains chitin and AF-647 phalloidin (red), F-actin. The xy planes move through increasing z depth, showing cuticular disks and F-actin mesh-like structures at different z positions of the scale cell.

**Supplementary Movie 2.** Planar sections through the same dataset (67% development) as depicted in Fig. S9. AF-555 WGA (green) stains chitin and AF-647 phalloidin (red), F-actin. Scale cells are relatively flatter at this stage

**Supplementary Movie 3.** Planar sections through the same dataset (76% development) as depicted in Fig. S10. AF-555 WGA (green) stains chitin and AF-647 phalloidin (red), F-actin. Cuticular disks have become extruded into hollow tubes and are enmeshed by tulip bulb-like F-actin structures.

**Supplementary Movie 4.** Planar sections through the same dataset as depicted in Figs. 5C and D, for *P. palinurus*. FITC WGA (green) stains chitin and TRITC phalloidin (red), F-actin. A concave whorl-like network of F-actin closely follow and underlie the cuticular dimples, in between ridges.

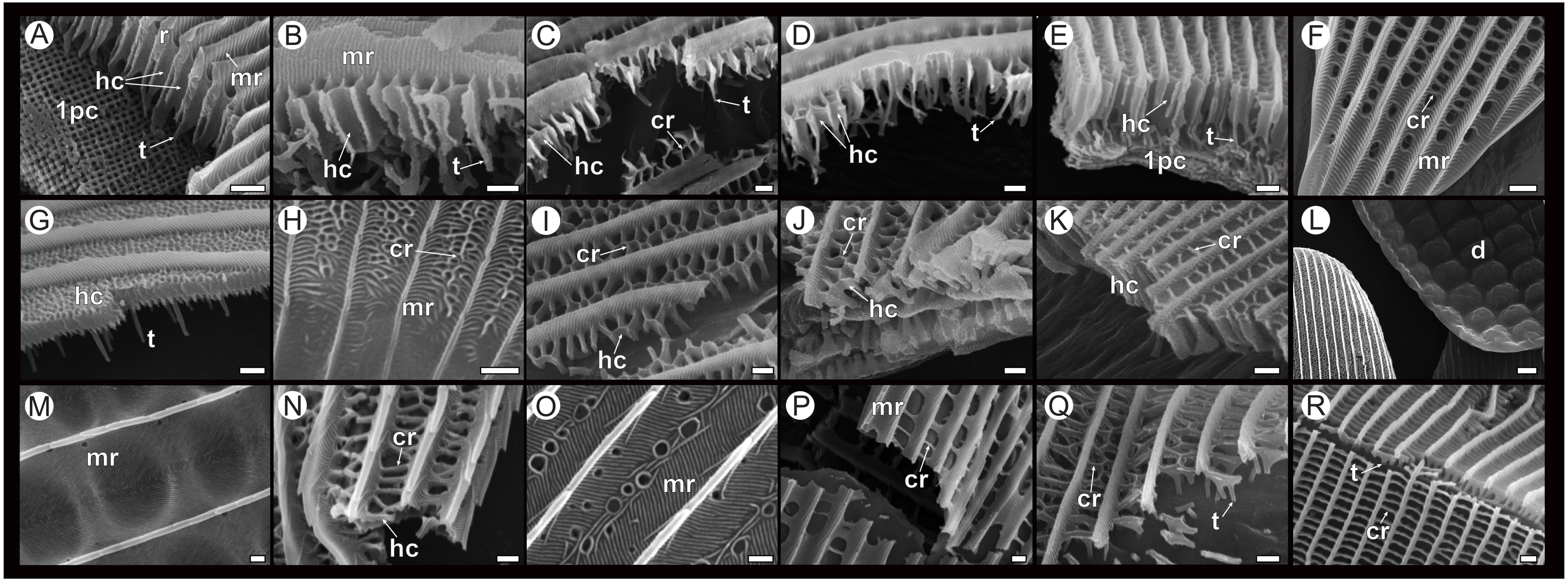

A
r
hc
mr
1pc
t
B
mr
hc
t
C
t
cr
hc
D
hc
t
E
hc
t
1pc
F
cr
mr
G
hc
t
H
cr
mr
I
cr
hc
J
cr
hc
K
cr
hc
L
d
M
mr
N
cr
hc
O
mr
P
mr
cr
Q
cr
t
R
t
cr

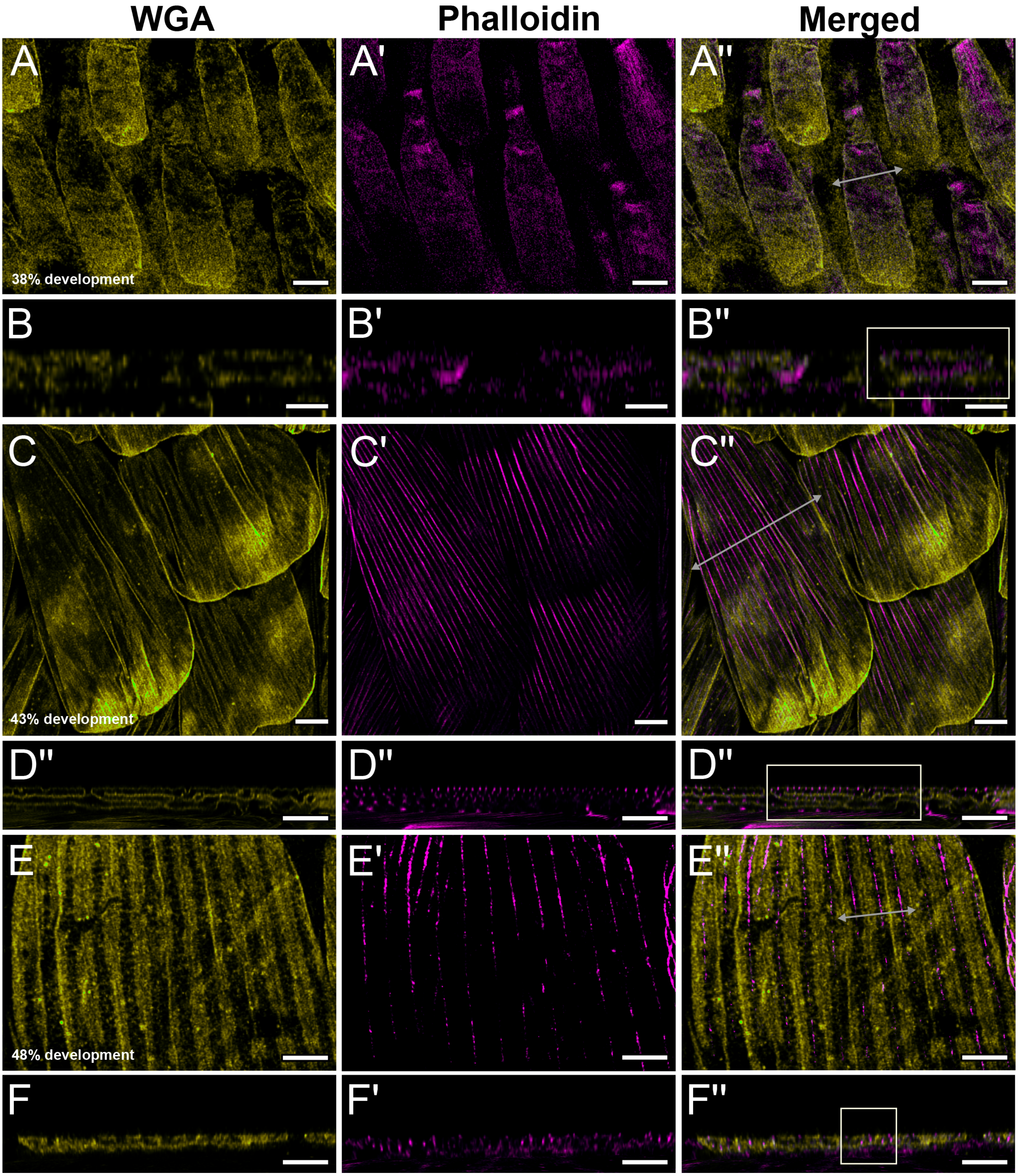
WGA
Phalloidin
Merged
A
A'
A''
38% development
B
B'
B''
C
C'
C''
43% development
D''
D''
D''
E
E'
E''
48% development
F
F'
F''

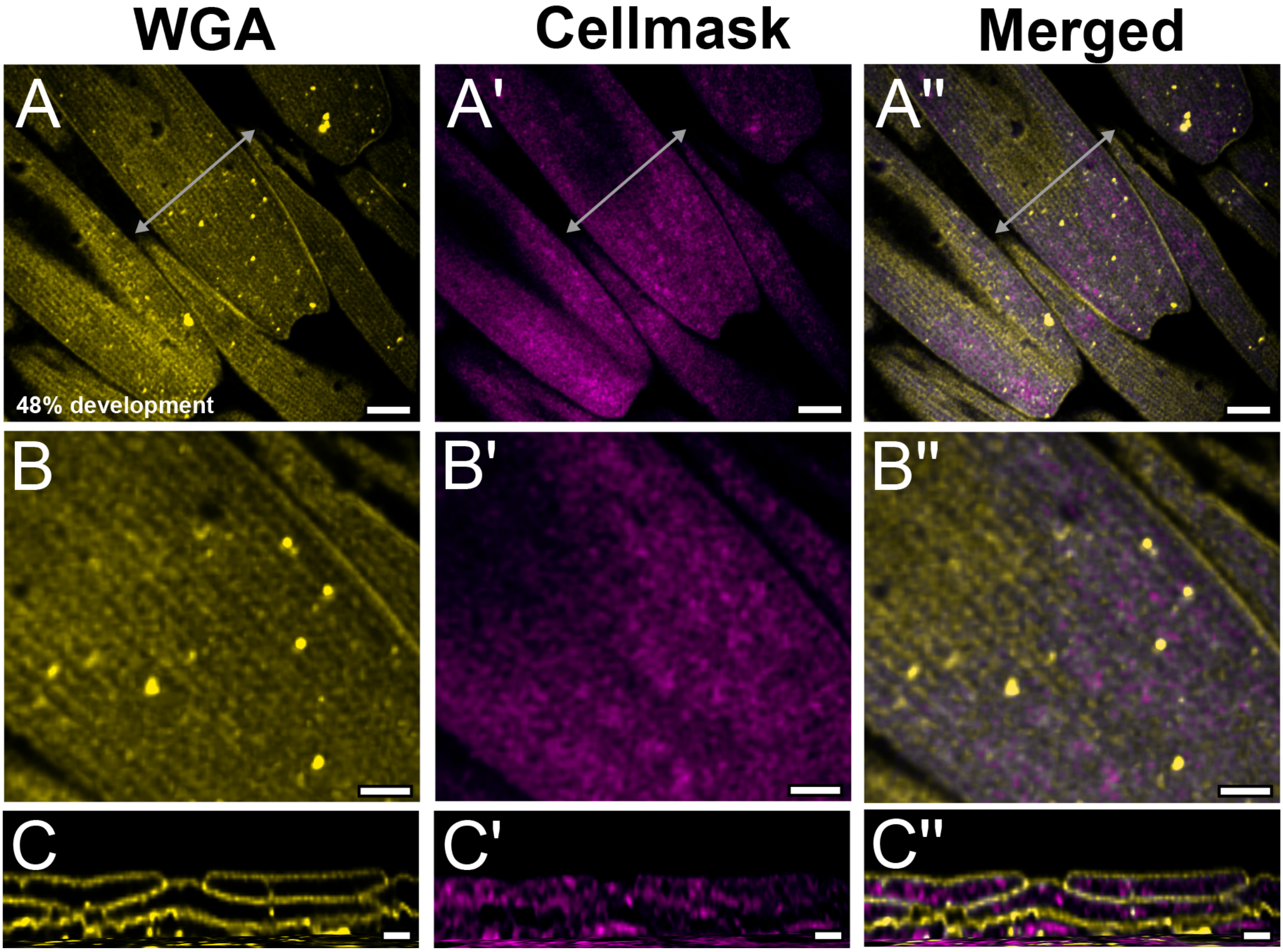
WGA
Cellmask
Merged
A
A'
A''
48% development
B
B'
B''
C
C'
C''

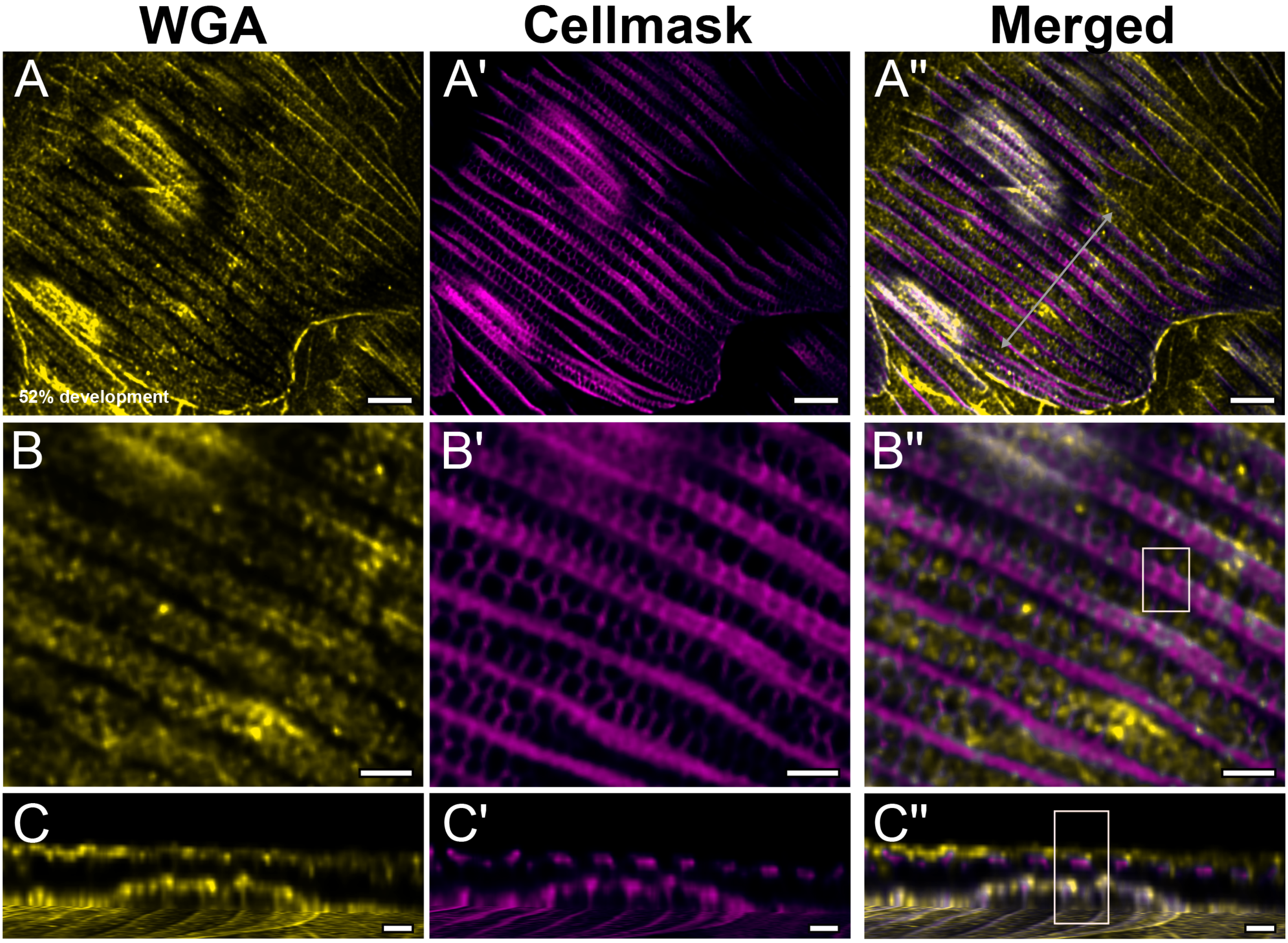
WGA
Cellmask
Merged
A
A'
A''
B
B'
B''
C
C'
C''
52% development

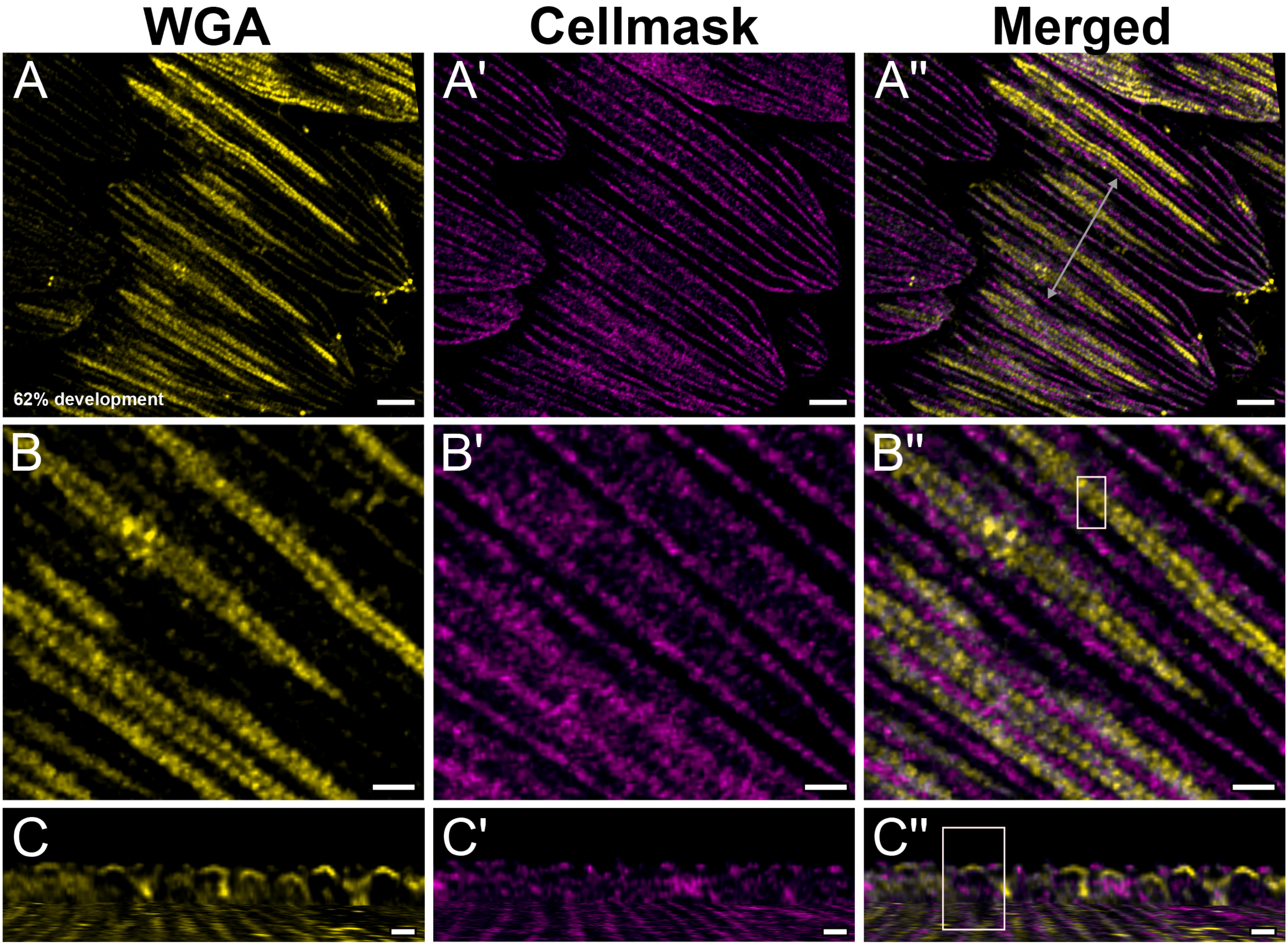
WGA
Cellmask
Merged
A
A'
A''
62% development
B
B'
B''
C
C'
C''

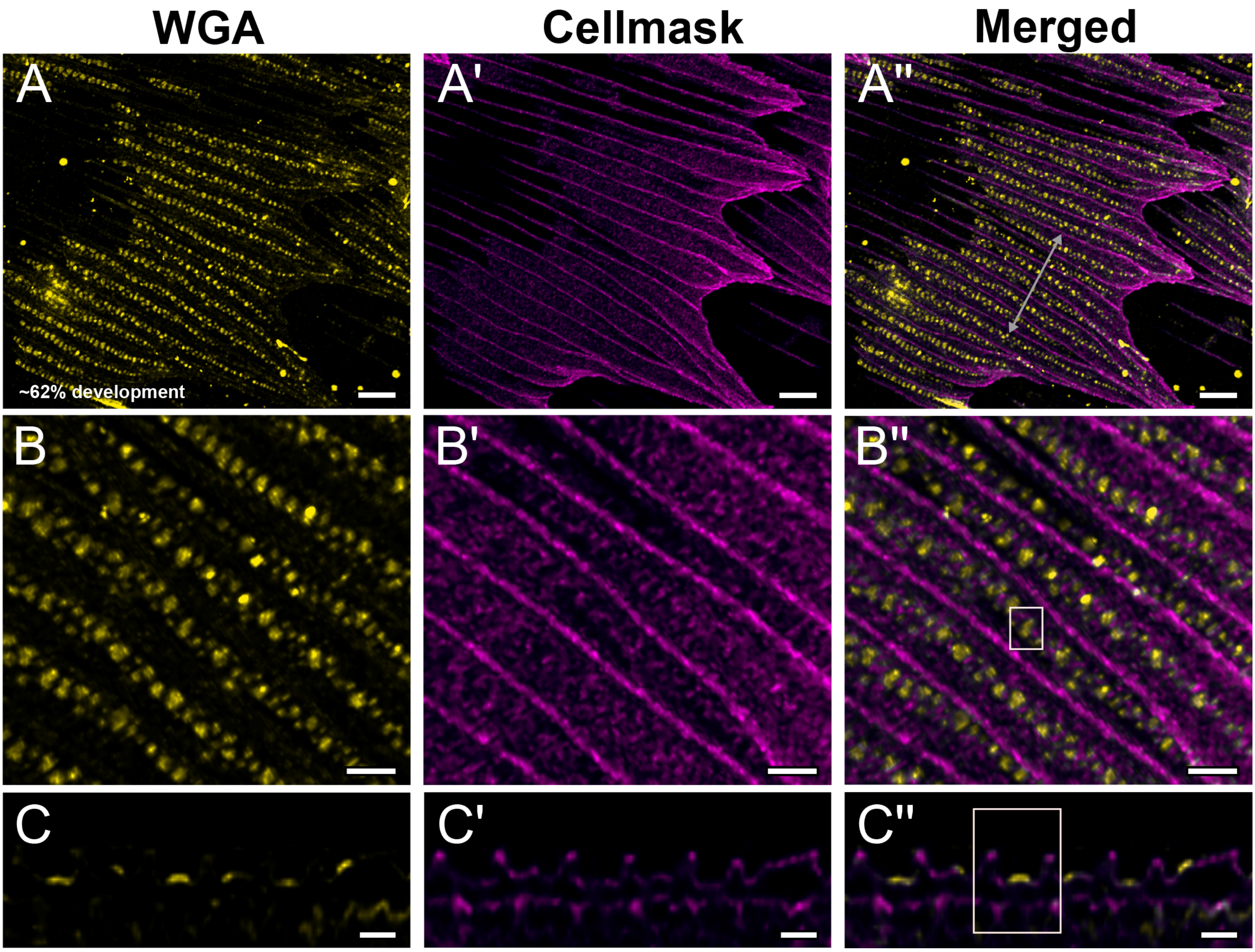

WGA
Cellmask
Merged
A
A'
A''
~62% development
B
B'
B''
C
C'
C''

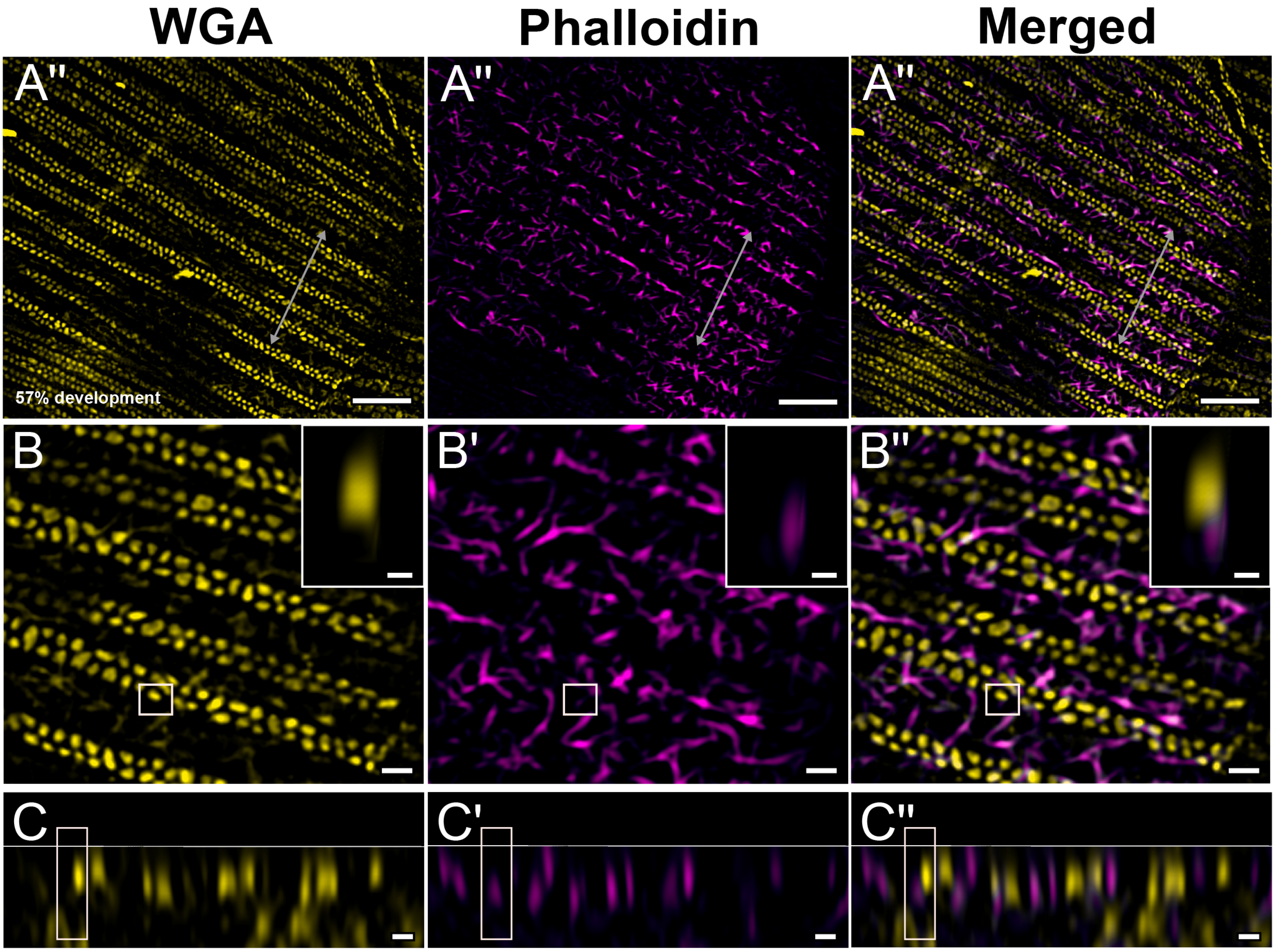

WGA
Phalloidin
Merged
A''
A''
A''
57% development
B
B'
B''
C
C'
C''

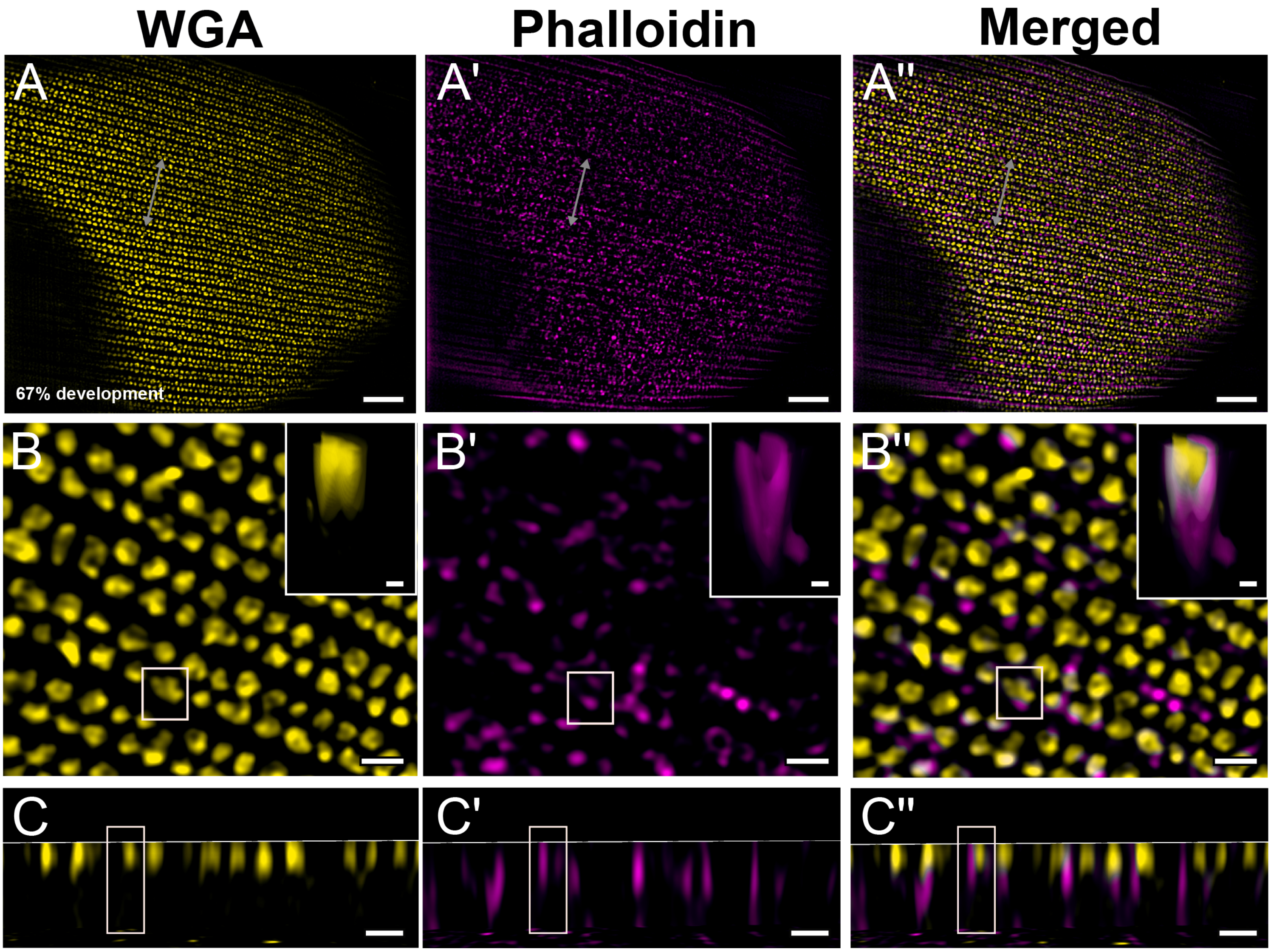

WGA
Phalloidin
Merged
A
A'
A''
67% development
B
B'
B''
C
C'
C''

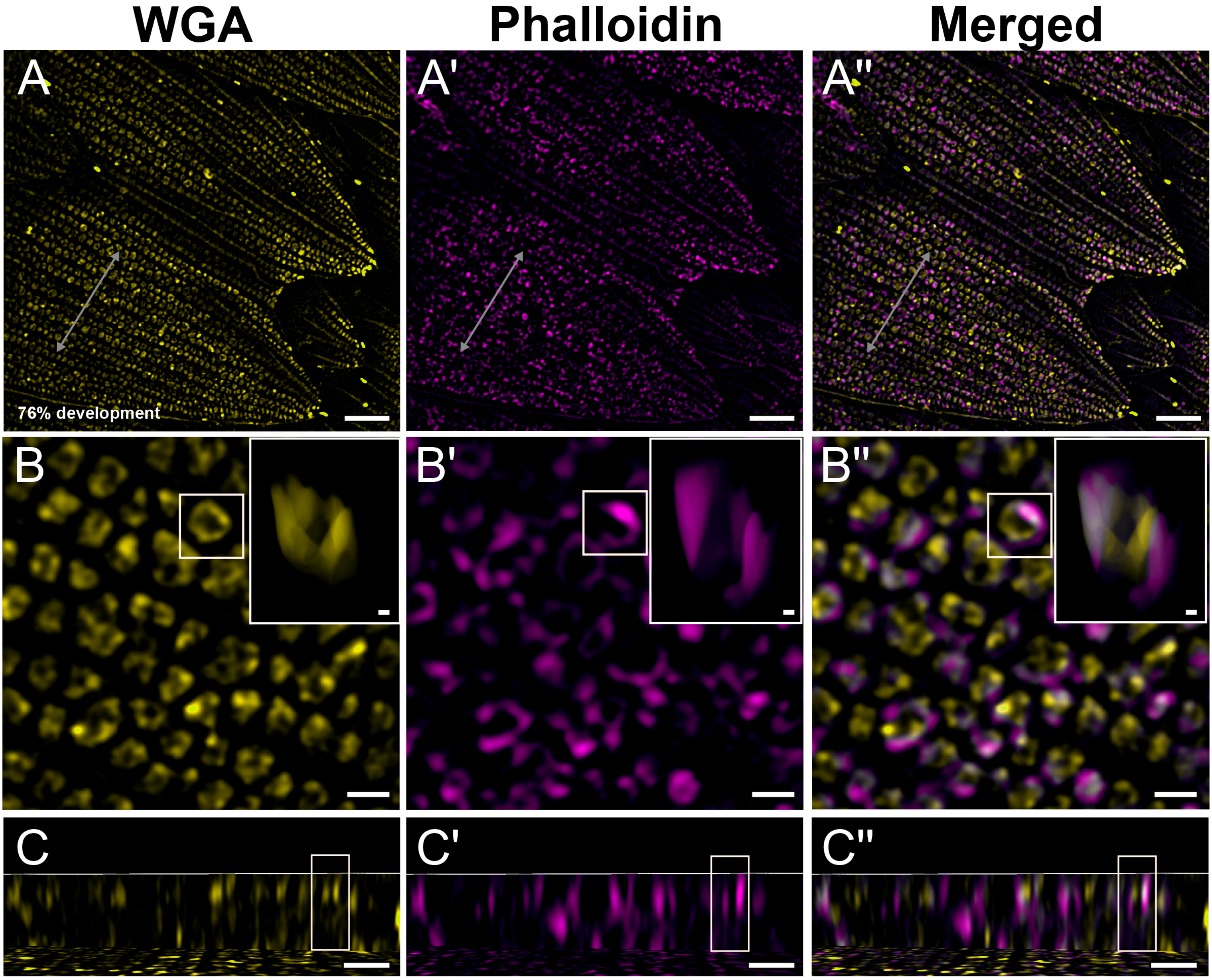
WGA
Phalloidin
Merged
A
A'
A''
76% development
B
B'
B''
C
C'
C''

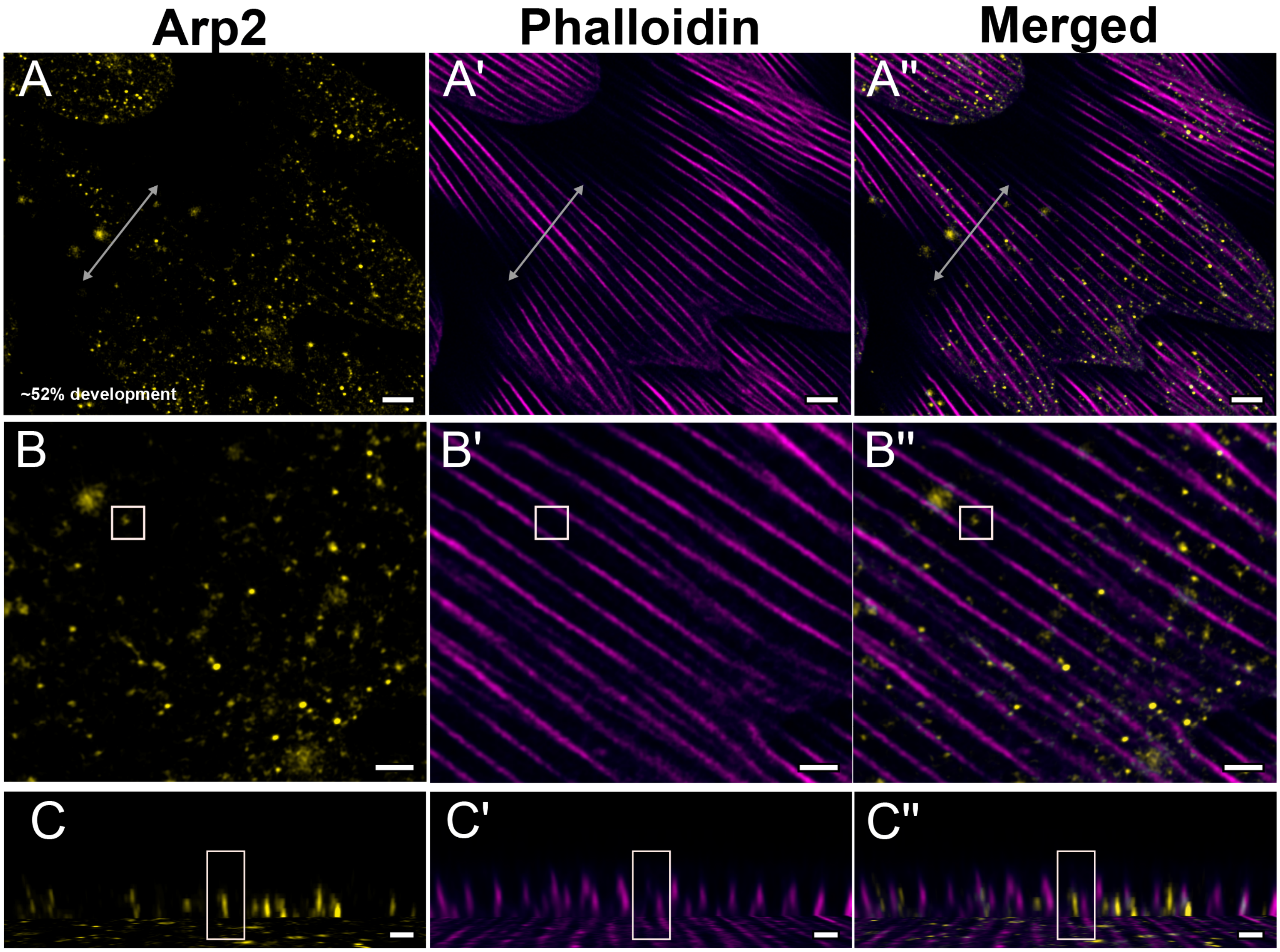

Arp2
Phalloidin
Merged
A
A'
A''
~52% development
B
B'
B''
C
C'
C''

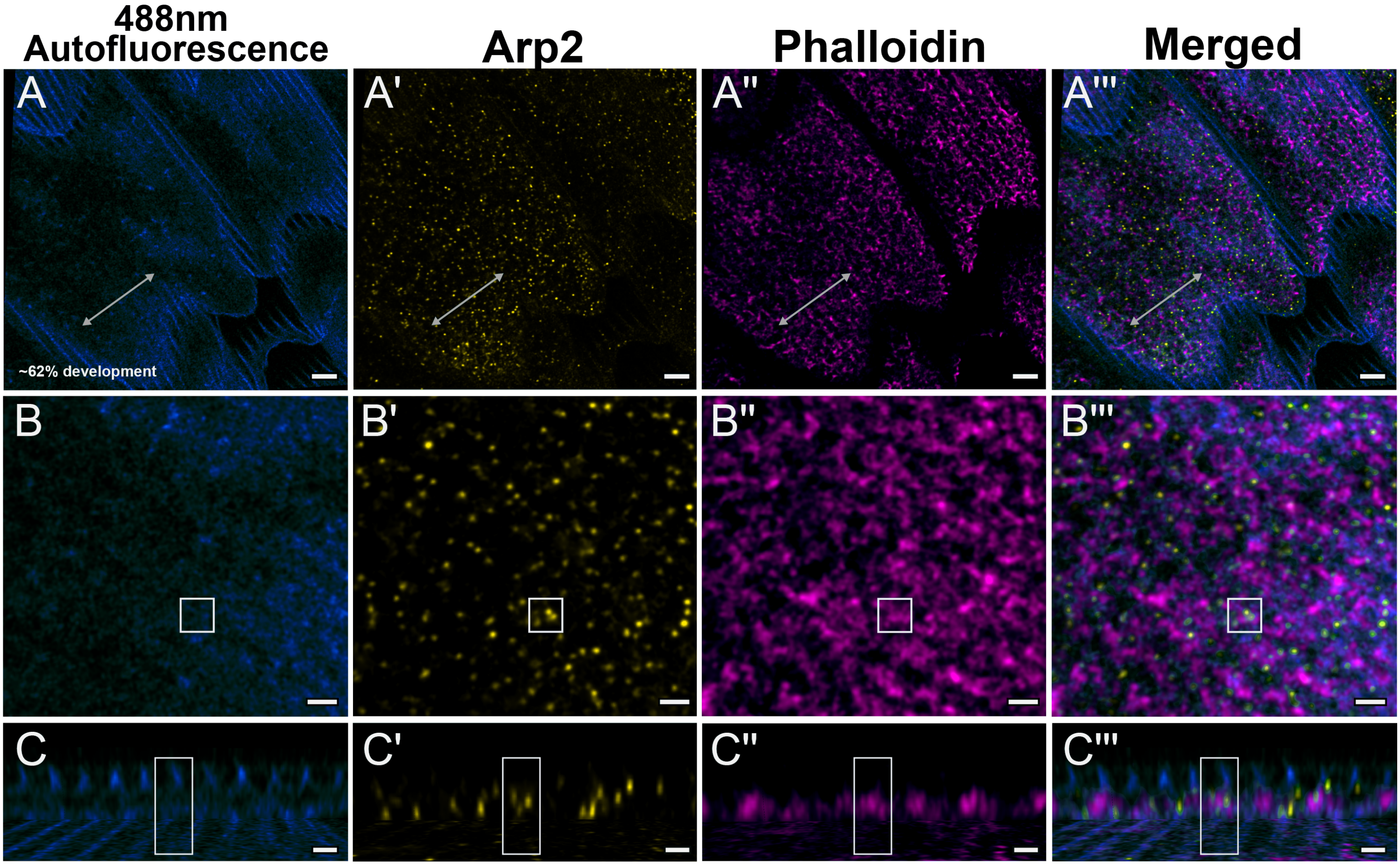

488nm Autofluorescence
Arp2
Phalloidin
Merged
A
A'
A''
A'''
~62% development
B
B'
B''
B'''
C
C'
C''
C'''

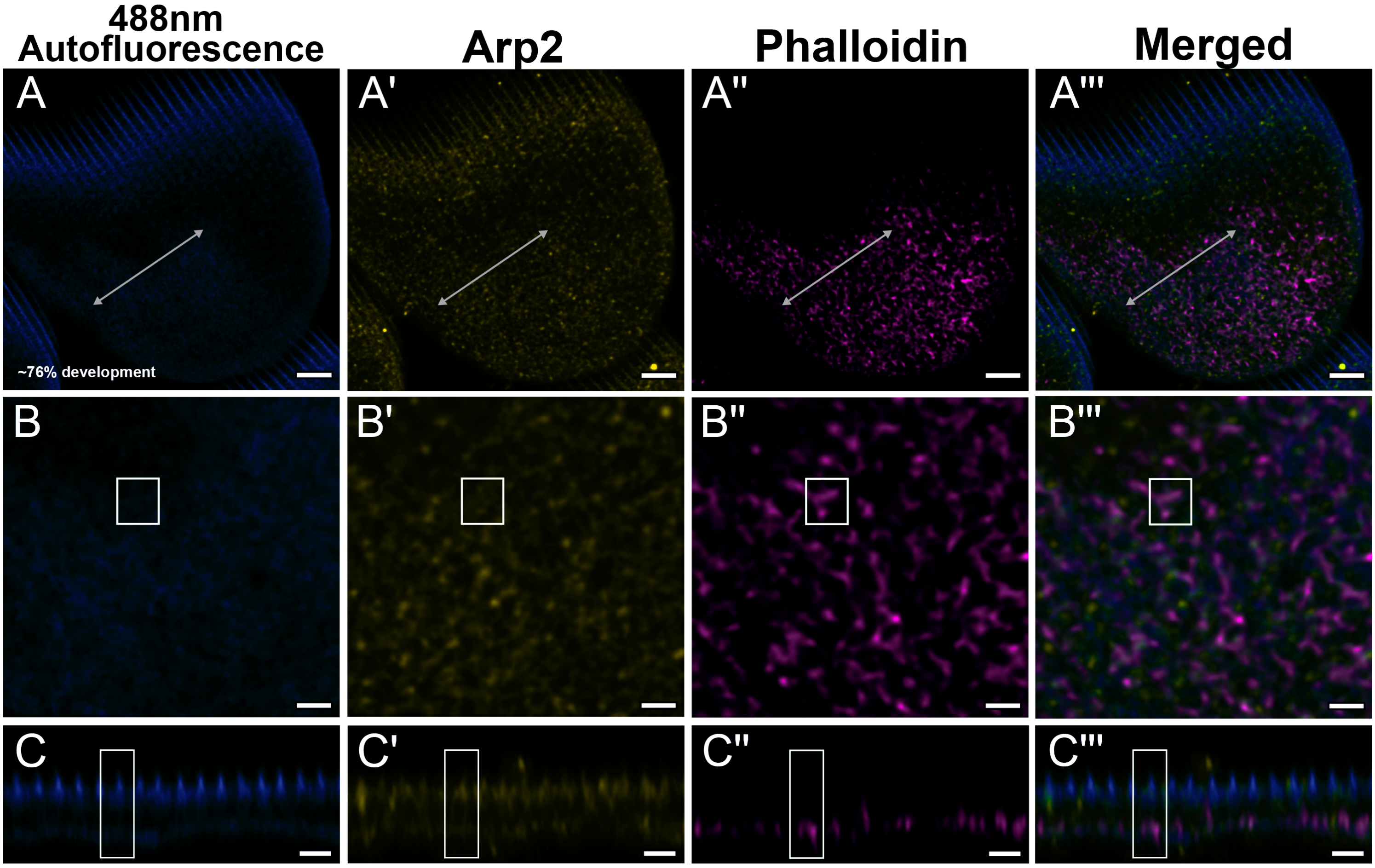
488nm Autofluorescence
Arp2
Phalloidin
Merged
A
A'
A''
A'''
~76% development
B
B'
B''
B'''
C
C'
C''
C'''

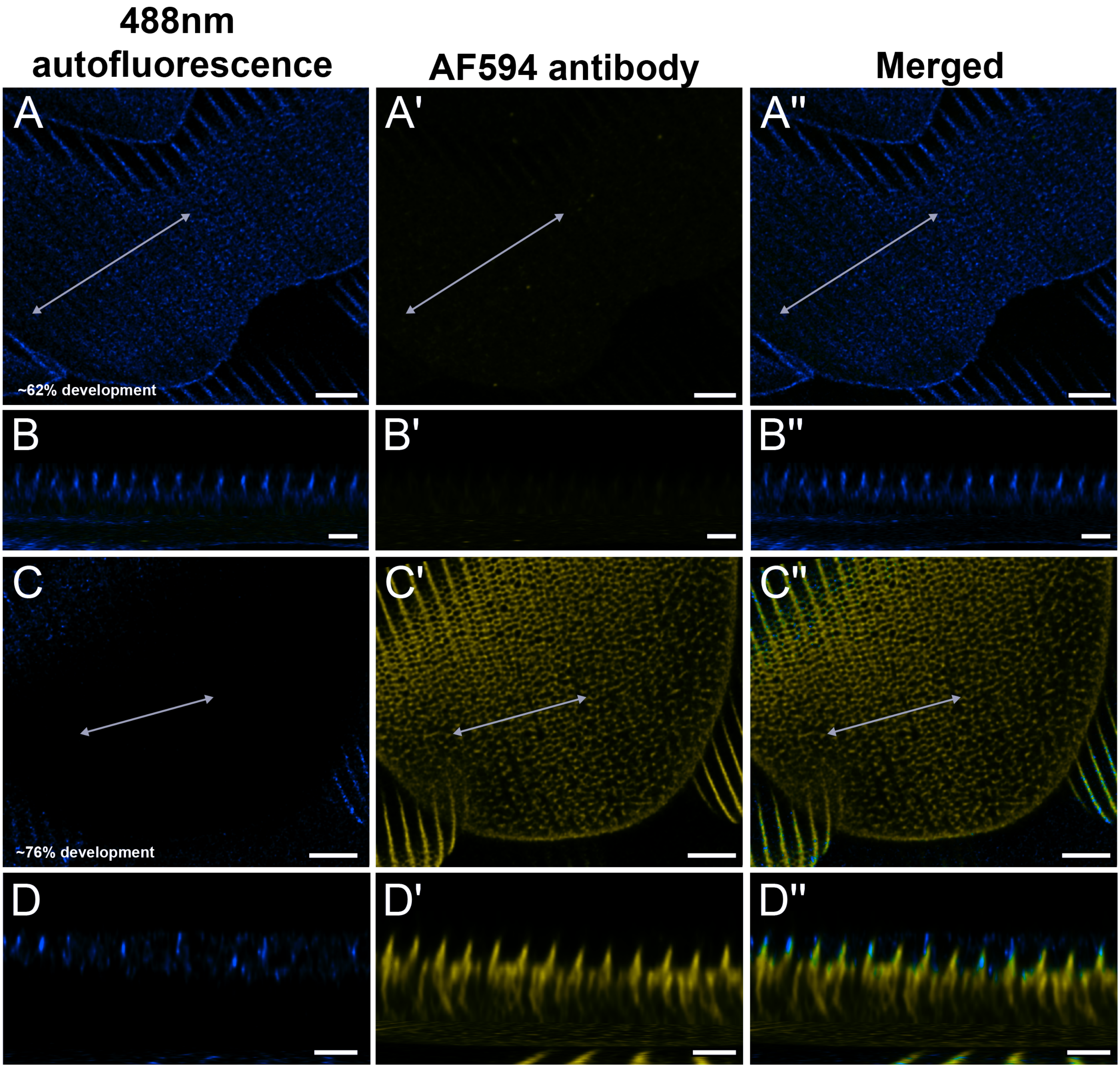

488nm autofluorescence
AF594 antibody
Merged
A
A'
A''
~62% development
B
B'
B''
C
C'
C''
~76% development
D
D'
D''

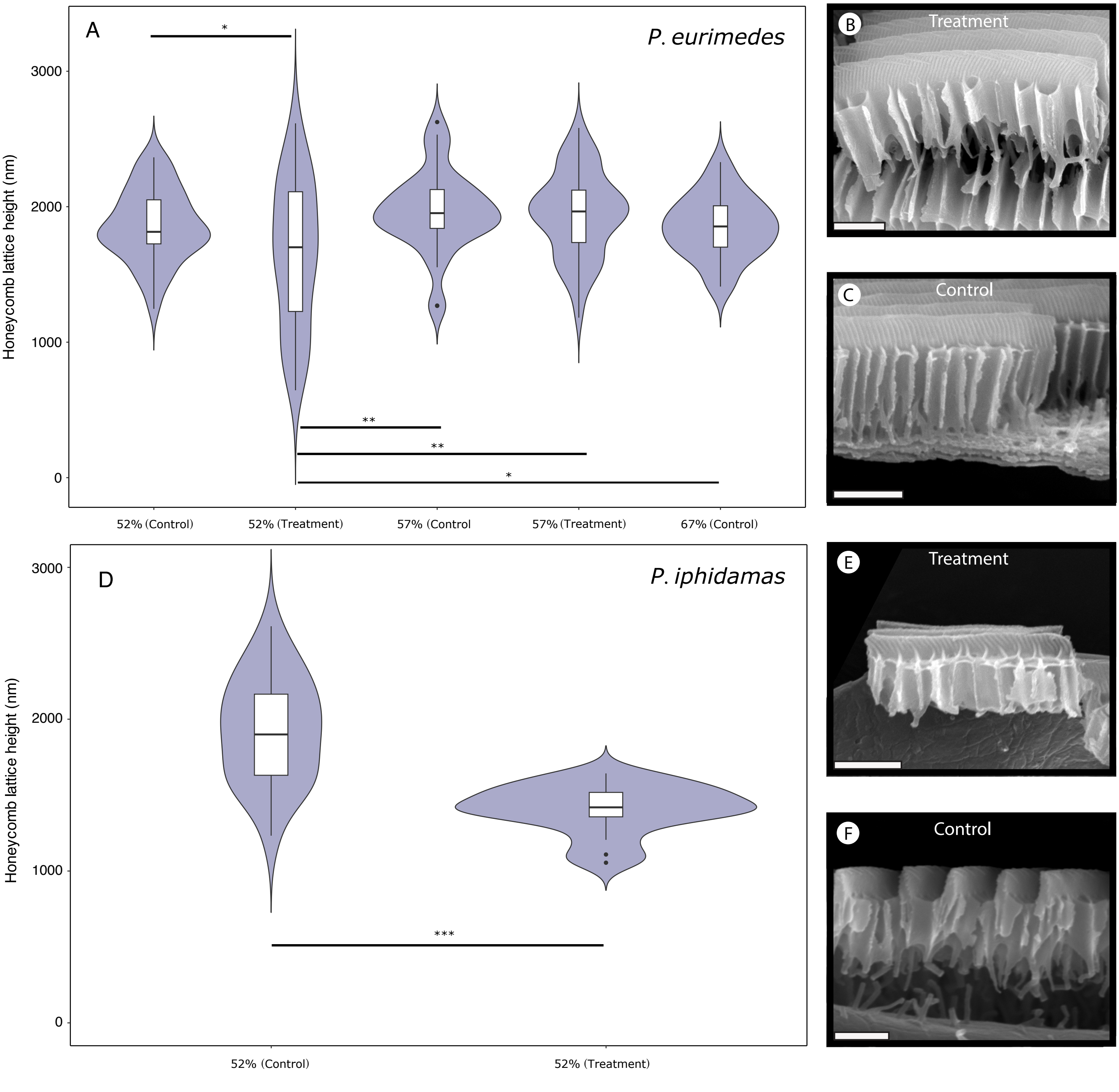

A
P. eurimedes
Honeycomb lattice height (nm)
3000
2000
1000
0
*
**
**
*
52% (Control)
52% (Treatment)
57% (Control
57% (Treatment)
67% (Control)
B
Treatment
C
Control
D
P. iphidamas
Honeycomb lattice height (nm)
3000
2000
1000
0
***
52% (Control)
52% (Treatment)
E
Treatment
F
Control

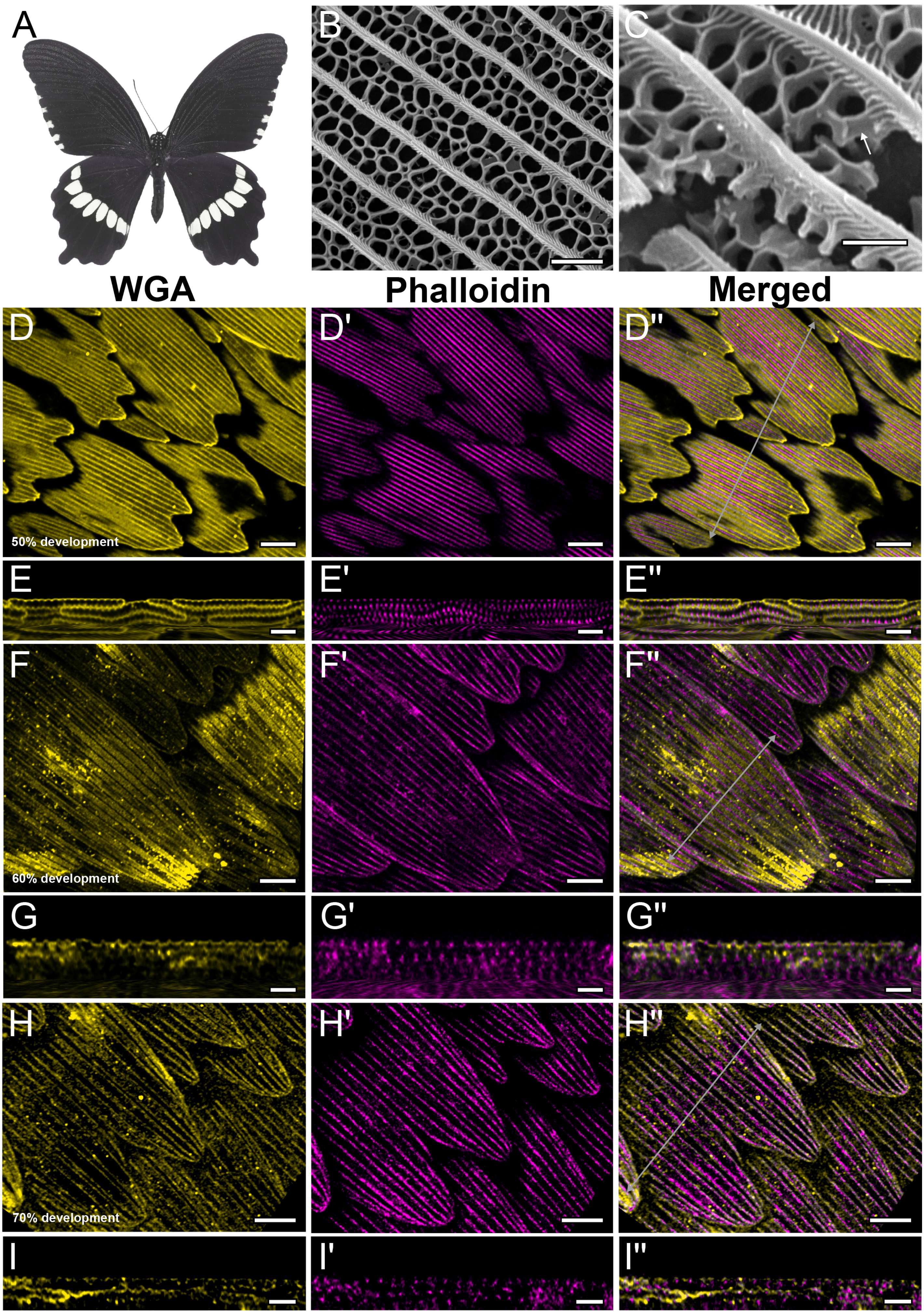
A
B
C
WGA
Phalloidin
Merged
D
D'
D''
50% development
E
E'
E''
F
F'
F''
60% development
G
G'
G''
H
H'
H''
70% development
I
I'
I''

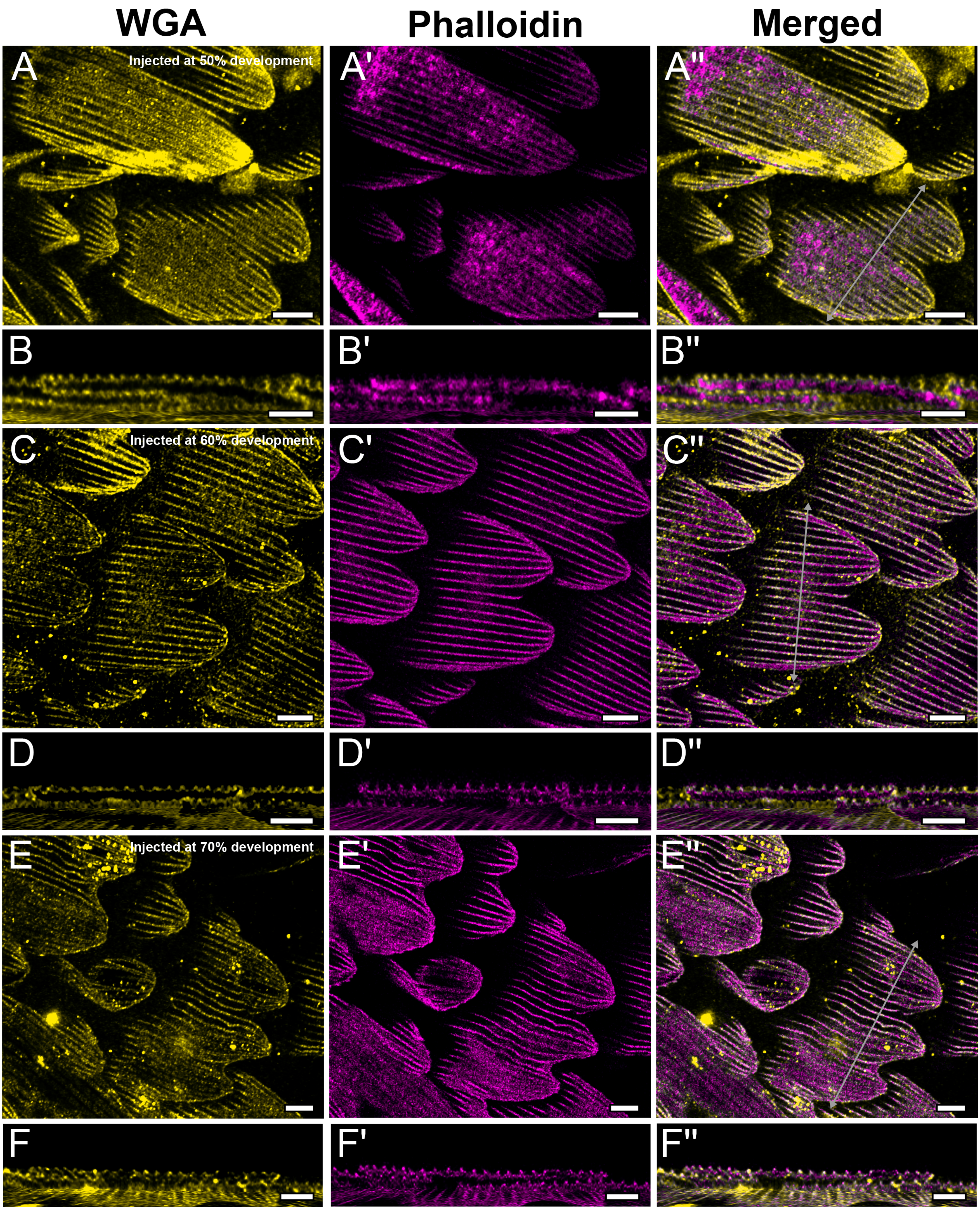
WGA
Phalloidin
Merged
A
Injected at 50% development
A'
A''
B
B'
B''
C
Injected at 60% development
C'
C''
D
D'
D''
E
Injected at 70% development
E'
E''
F
F'
F''

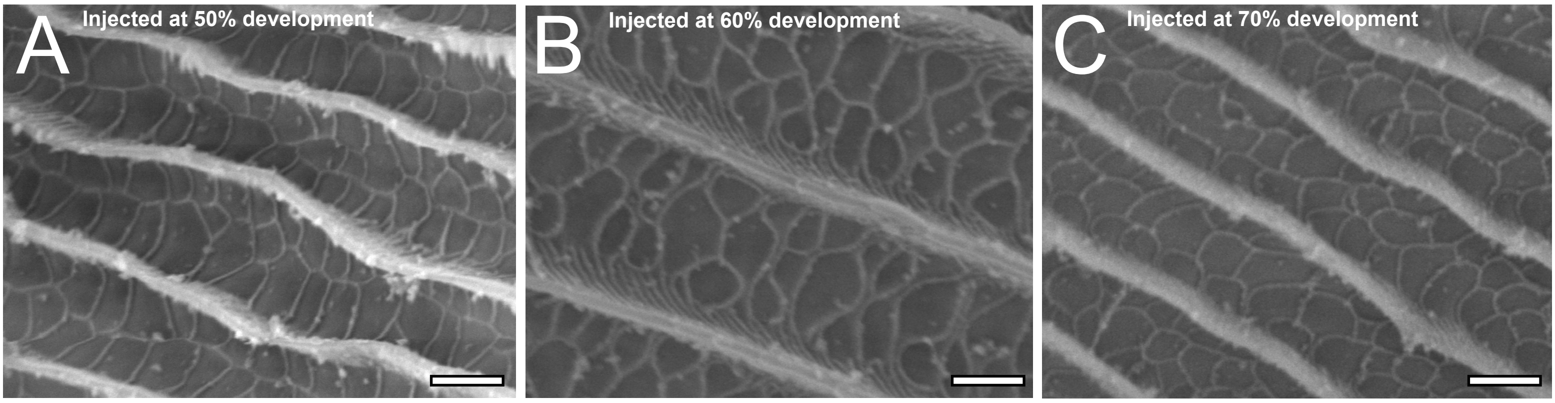
A
Injected at 50% development
B
Injected at 60% development
C
Injected at 70% development

**Table S1.** Details of CK-666 inhibition experiments performed in different batches of purchased pupae and their eventual fates.

| Species | Sample no. | Sex | Treatment | Pupation stage when injected | Fate |
|---|---|---|---|---|---|
| *P. eurimedes* | 001 | F | - | - | Dead |
| *P. eurimedes* | 002 | M | 100uM CK-666 | 52% | Emerged* |
| *P. eurimedes* | 003 | M | DMSO control | 52% | Emerged* |
| *P. eurimedes* | 004 | M | 100uM CK-666 | 57% | Emerged* |
| *P. eurimedes* | 005 | M | 100uM CK-666 | 57% | Emerged* |
| *P. eurimedes* | 006 | M | DMSO control | 57% | Emerged* |
| *P. eurimedes* | 007 | M | 100uM CK-666 | 67% | Dead |
| *P. eurimedes* | 008 | M | 100uM CK-666 | 67% | Dead |
| *P. eurimedes* | 009 | M | DMSO control | 67% | Emerged* |
| *P. iphidamas* | 001 | M | 100uM CK-666 | 52% | Emerged* |
| *P. iphidamas* | 002 | F | 100uM CK-666 | 52% | Emerged* |
| *P. iphidamas* | 003 | M | DMSO control | 52% | Emerged* |
| *P. iphidamas* | 004 | M | 100uM CK-666 | 67% | Dead |
| *P. iphidamas* | 005 | M | 100uM CK-666 | 67% | Dead |
| *P. iphidamas* | 006 | M | DMSO control | 67% | Dead |
| *P. polytes* | 001 | M | 10uM CK-666 | 40% | Dissected |
| *P. polytes* | 002 | F | 10uM CK-666 | 40% | Dissected |
| *P. polytes* | 003 | F | 10uM CK-666 | 40% | Dissected |
| *P. polytes* | 004 | F | DMSO control | 40% | Emerged |
| *P. polytes* | 005 | M | 100uM CK-666 | 40% | Dissected |

| | | | | | |
|---|---|---|---|---|---|
| *P. polytes* | 006 | F | 100uM CK-666 | 40% | Dissected |
| *P. polytes* | 007 | F | 100uM CK-666 | 40% | Dissected |
| *P. polytes* | 008 | F | DMSO control | 40% | Dissected |
| *P. polytes* | 009 | F | 10uM CK-666 | 50% | Dissected |
| *P. polytes* | 010 | M | 10uM CK-666 | 50% | Dissected |
| *P. polytes* | 011 | F | 10uM CK-666 | 50% | Dissected |
| *P. polytes* | 012 | M | DMSO control | 50% | Emerged |
| *P. polytes* | 013 | M | 100uM CK-666 | 50% | Dissected |
| *P. polytes* | 014 | F | 100uM CK-666 | 50% | Dissected |
| *P. polytes* | 015 | F | 100uM CK-666 | 50% | Dissected |
| *P. polytes* | 016 | M | DMSO control | 50% | Dissected |
| *P. polytes* | 017 | M | 10uM CK-666 | 60% | Dissected |
| *P. polytes* | 018 | F | 10uM CK-666 | 60% | Dissected |
| *P. polytes* | 019 | M | 10uM CK-666 | 60% | Dissected |
| *P. polytes* | 020 | M | DMSO control | 60% | Dissected |
| *P. polytes* | 021 | F | 100uM CK-666 | 60% | Dissected |
| *P. polytes* | 022 | F | 100uM CK-666 | 60% | Dissected |
| *P. polytes* | 023 | M | 100uM CK-666 | 60% | Dissected |
| *P. polytes* | 024 | F | 100uM CK-666 | 60% | Dissected |
| *P. polytes* | 025 | M | DMSO control | 60% | Dissected |
| *P. polytes* | 026 | F | 10uM CK-666 | 70% | Dissected |
| *P. polytes* | 027 | F | 10uM CK-666 | 70% | Dissected |
| *P. polytes* | 028 | F | DMSO control | 70% | Dissected |
| *P. polytes* | 029 | F | 100uM CK-666 | 70% | Dissected |
| *P. polytes* | 030 | F | 100uM CK-666 | 70% | Emerged |

| | | | | | |
|---|---|---|---|---|---|
| *P. polytes* | 031 | F | - | - | Dead |
| *P. polytes* | 032 | F | 100uM CK-666 | 70% | Dissected |
| *P. polytes* | 033 | M | DMSO control | 70% | Dissected |
| *P. polytes* | 034 | M | 10uM CK-666 | 80% | Dissected |
| *P. polytes* | 035 | F | 10uM CK-666 | 80% | Dissected |
| *P. polytes* | 036 | F | DMSO control | 80% | Dissected |
| *P. polytes* | 037 | F | 100uM CK-666 | 80% | Dissected |
| *P. polytes* | 038 | F | 100uM CK-666 | 80% | Dissected |
| *P. polytes* | 039 | F | 100uM CK-666 | 80% | Dissected |
| *P. polytes* | 040 | F | DMSO control | 80% | Dissected |

*All *Parides eurimedes* and *P. iphidamas* were allowed to develop and emerge as adults and then dissected, unlike *Papilio polytes* pupae (see Methods).
